# A BTB extension and ion-binding domain contribute to the pentameric structure and TFAP2A binding of KCTD1

**DOI:** 10.1101/2024.06.14.599093

**Authors:** Daniel M. Pinkas, Joshua C. Bufton, Alice E. Hunt, Charlotte E. Manning, William Richardson, Alex N. Bullock

**Affiliations:** Centre for Medicines Discovery, Nuffield Department of Medicine Research Building, University of Oxford, Roosevelt Drive, Oxford, OX3 7FZ, UK; Department of Biological Sciences, Universidad Loyola Andalucía, Seville, Spain

**Keywords:** BTB domain, X-ray crystallography, ion channel, scalp-ear-nipple syndrome, pentamer, Cullin-RING ligase, Cullin3, TFAP2, KCTD, beta-propeller

## Abstract

KCTD family proteins typically assemble into Cullin-RING E3 ligases. KCTD1 is an atypical member that functions instead as a transcriptional repressor. Mutations in KCTD1 cause developmental abnormalities and kidney fibrosis in scalp-ear-nipple syndrome. Here, we present unexpected mechanistic insights from the structure of full-length KCTD1. Disease-causing mutation P20S maps to an unrecognized extension of the BTB domain that contributes both to its pentameric structure and TFAP2A binding. The C-terminal domain (CTD) shares its fold and pentameric assembly with the protein GFRP despite lacking discernible sequence similarity. Most surprisingly, the KCTD1 CTD establishes a central channel occupied by alternating sodium and iodide ions that restrict TFAP2A dissociation. The elucidation of the structure redefines the KCTD family BTB domain fold and identifies an unexpected ion pore for future study of KCTD1’s function in the ectoderm, neural crest and kidney.

## INTRODUCTION

The 25 members of the KCTD family form a subgroup of BTB domain-containing proteins that commonly assemble as cullin3-dependent E3 ligases and use a variable C-terminal domain for substrate recognition^1-4^. Structural studies have been central to their characterization. The prototypical KCTD5 was the first member to have its protein structure elucidated and remains the only family member for which the available structures span both domains^5-7^. This protein is observed to act as an off-switch for GPCR signalling through the ubiquitin-mediated degradation of G-protein βγ subunits^6,8,9^. Its BTB domain displays significant structural conservation with the potassium (K^+^) channel tetramerization domain (T1), from which the KCTD family derives its name, but adopts a closed pentameric assembly that binds to cullin3 with 5:5 stoichiometry^5,7^. The associated pentameric CTD of KCTD5 resembles a beta-propeller-like structure that can similarly assemble as a 5:5 complex with Gβγ subunits, further highlighting the importance of oligomerization and co-operative binding for KCTD family function^6,7^.

Not all KCTD family proteins demonstrate binding to cullin3^3,4,10^. Structural and biophysical studies of the Clade F proteins KCTD8, KCTD12, and KCTD16 have provided insights into their function instead as auxiliary subunits of GABA_B_ receptors. Tethered to GABA_B2_ by their BTB domains, the CTDs of these KCTD proteins bind to Gβγ subunits released from Gα, resulting in rapid deactivation of downstream GIRK channel activity^11-14^. Remarkably, the structure of the 5:5 KCTD12-Gβγ complex reveals an expanded CTD domain (the H1 domain) that exhibits an entirely distinct protein-protein interface to the equivalent KCTD5 complex^11^. Indeed, recent work suggests that Gβγ interaction may be a common trait within the different KCTD family subgroups despite their diverged structural mechanisms^9^.

The clade A proteins KCTD1 and KCTD15 form a further distinct, but important subgroup with no binding to cullin3 and no apparent role in G-protein regulation^3,10^. These proteins instead repress the transactivation domain of the transcription factor TFAP2A (AP-2α)^15,16^. Importantly, disease-causing mutations in KCTD1, KCTD15 and TFAP2A that abrogate this function are associated with craniofacial abnormalities and cutis aplasia^15,17-20^. Missense substitutions in the BTB domain of KCTD1 cause scalp-ear-nipple (SEN) syndrome, an autosomal dominant condition characterized by facial dysmorphism and ectodermal abnormalities, including sparse hair and the absence of nipples, as well as progressive renal fibrosis^21,22^. Mutant KCTD1 proteins display reduced TFAP2A binding and thermostability resulting in a propensity to form amyloid-like aggregates and dominant negative loss of function in mixed models of wild type and mutant^15,18,19^. Their effects are partly understood from structures of the truncated BTB domain^3^, with the notable exception of the P20S mutation which falls outside the known domains of the KCTD family.

Here, we analyzed the structure, stability and interactions of full-length KCTD1 to better understand the functionally important N-terminus, the spatial arrangement of BTB-CTD domains and the previously uncharacterized CTD fold. We uncovered unexpected structural features across the length of KCTD1 that contribute to its pentameric assembly and TFAP2A interaction.

## RESULTS

### Structure of full-length KCTD1

To understand of the structural mechanisms of the multidomain KCTD1 protein, we purified both full length and truncated forms of the KCTD1 protein and performed crystallisation screening. Recombinant human KCTD1 was observed to crystallize in coarse screen conditions containing sodium chloride or sodium iodide. Fine screening revealed that high concentrations of sodium iodide were required for reproducible diffracting crystals. A 2.4 Å structure was initially obtained for KCTD1_Δ__N27_ (removing 27 N-terminal residues) and subsequently a 2.7 Å structure of the full-length protein (Figures S1 and 1, Table S1). The structures reveal that KCTD1 forms a closed homopentamer, as found for full-length KCTD5. A short helical linker connects the BTB and CTD domains, which share nearly coaxial C5 symmetry axes. The CTDs are twisted around that axis by a full pentameric subunit with respect to the BTB domains.

Only 3 of the 101 residues in the CTD of KCTD1 are conserved in KCTD5. Yet, remarkably, the domains are found to be structurally related (RMSD = 2.2 Å over 50 Cα atoms), with KCTD1 exhibiting a large insertion with respect to the CTD of KCTD5, nearly doubling its size from 6 kDa to 11 kDa. While the KCTD5 CTD displays a relatively ‘open’ solvent-filled central channel, the CTD of KCTD1 exhibits a tightly restricted channel occupied by sodium and iodide ions sequestered from the crystallization solution (Figures S1 and 1).

### An extended BTB domain in KCTD1 is required for TFAP2 binding

A most unexpected and striking feature of the full-length KCTD1 structure was the oligomeric assembly of the BTB domain. Previous KCTD family structures were truncated to the globular BTB domain as in KCTD1_Δ__N27_ . The full-length KCTD1 structure reveals a hitherto unrecognized 11-residue N-terminal extension of the BTB domain that drapes across the neighbouring BTB domain to establish a ring of intersubunit interactions around the outside of the BTB pentamer (Figure 2A). Importantly, this packing provides a structural explanation for the disease-causing KCTD1 mutations P20S and G62D (and equivalent KCTD15 mutation G88D) that are known to destabilise the proteins and reduce TFAP2 binding (Figures 2A-B). The structure reveals that Pro20 falls within the N-terminal extension and inserts into a hydrophobic pocket in the neighbouring BTB to contact residues Leu45, Phe60, Gly62, Ile66 and Tyr75 in this domain. The mutations P20S and G62D would abrogate this hydrophobic packing and affect the backbone conformation. Consistent with this mutation, KCTD1_Δ__N27_ lacking the N-terminal extension displayed minimal binding to a TFAP2A peptide derived from the transactivation domain, whereas full-length KCTD1 exhibited submicromolar affinity further highlighting the importance of the N-terminal BTB extension for KCTD1 function (Figures 2C-D). The full-length KCTD1 structure was obtained from a crystallisation drop containing TFAP2A peptide with the hope to determine its co-structure. However, the resulting electron density maps provided no evidence for bound peptide and so the precise structural basis for its interaction remains to be elucidated.

**Figure 1.**
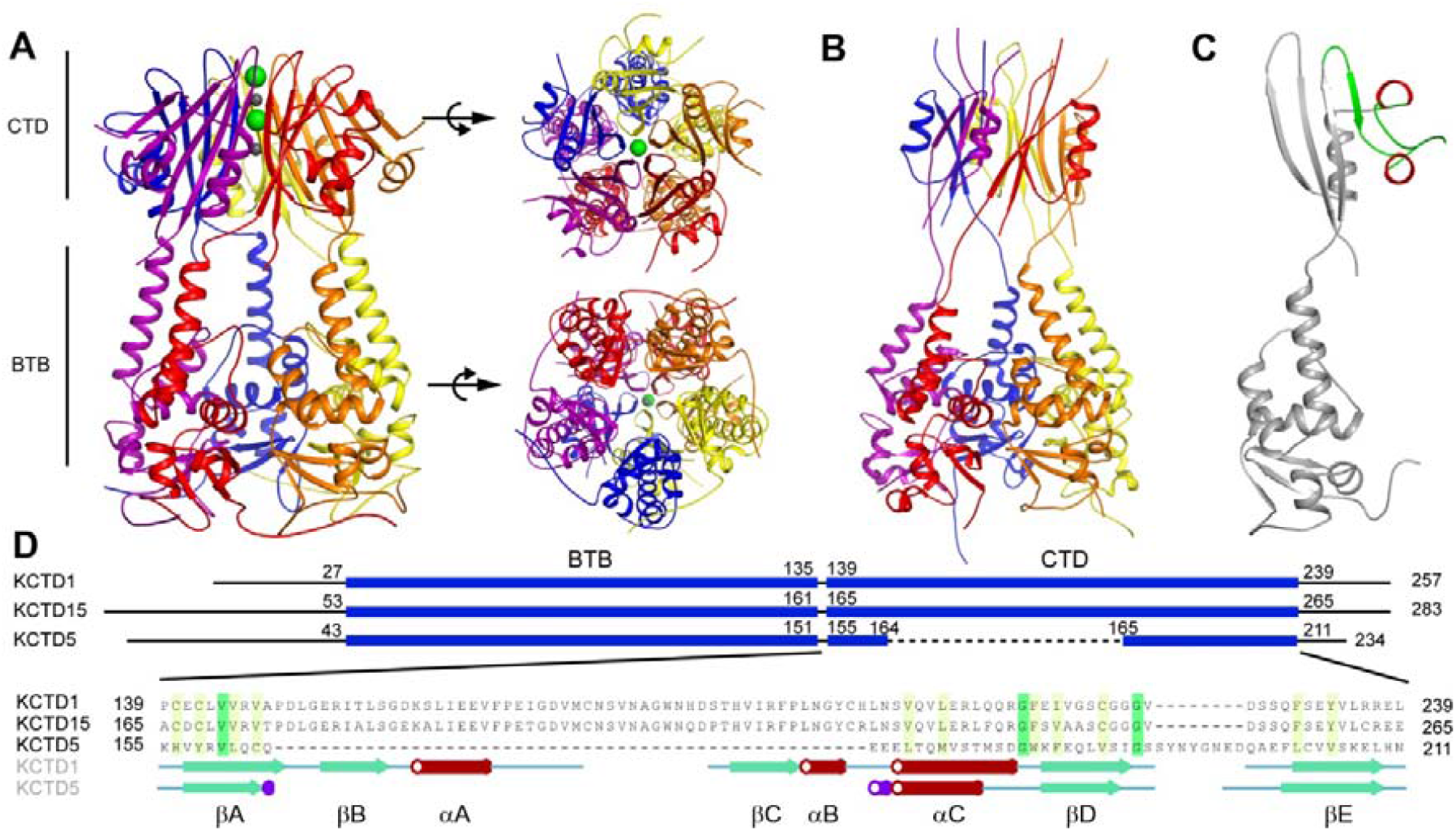
Structure of full-length KCTD1. (**A**) Ribbon diagram of the pentameric KCTD1 structure. Sodium and iodide ions are shown as gray and green spheres, respectively (**B**) Previously determined KCTD5 structure (PDB 3DRY). (**C**) Single chain of KCTD1. An insertion in the CTD not found in KCTD5 is highlighted in red (helix) and green (β-strand and coil). (**D**) Structure-based alignment of the CTDs of KCTD1, KCTD15 and KCTD5. See also Figure S1.

**Figure 2.**
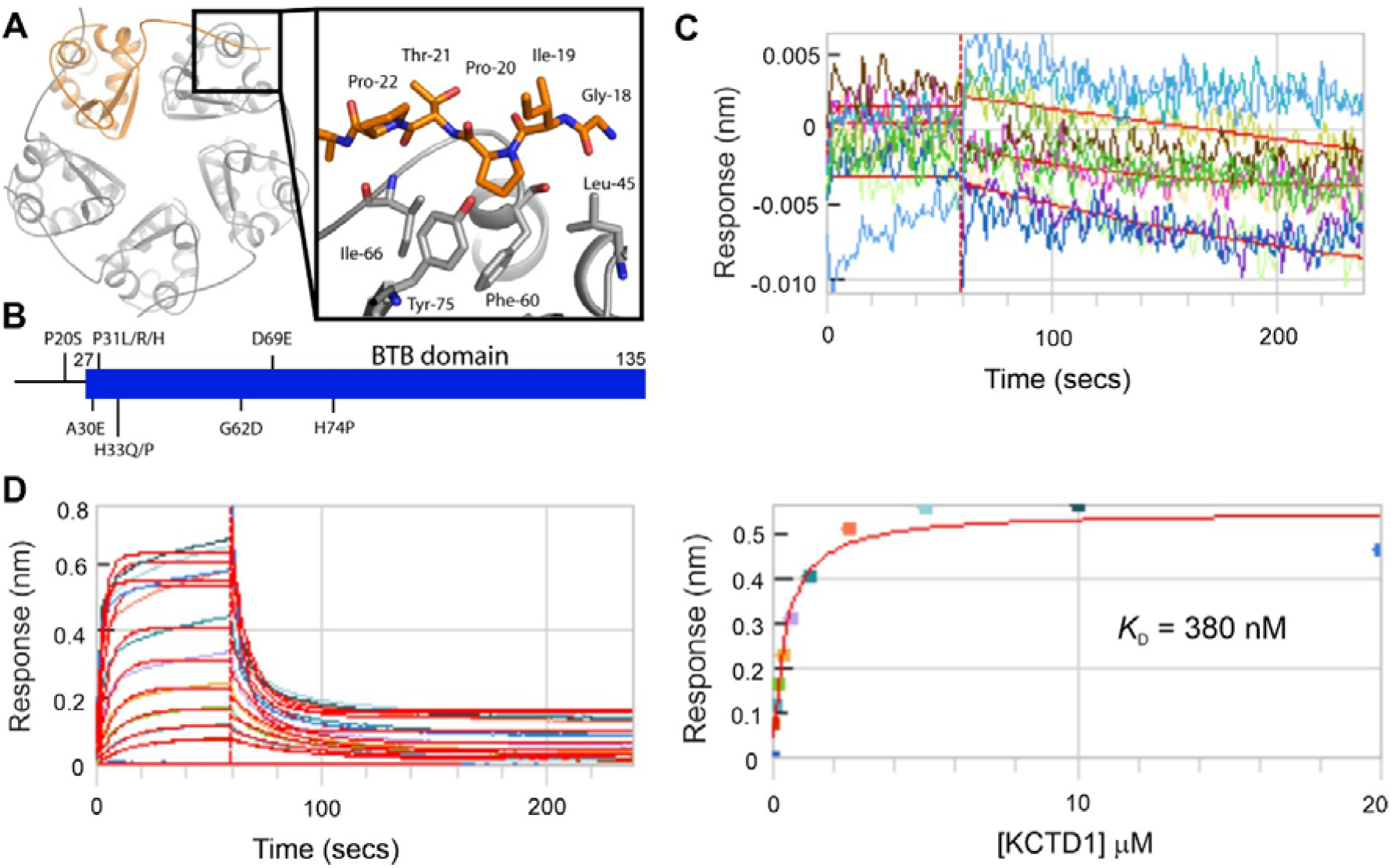
An extended BTB domain in KCTD1 is required for TFAP2 binding. (**A**) Pentameric BTB domain assembly highlighting one chain in orange. Inset shows interactions of the N-terminal BTB extension with the neighbouring subunit. (**B**) Locations of disease-causing missense mutations in KCTD1. (**C-D**) BLI measurements. Biotinylated TFAP2A peptide was immobilised on streptavidin-coated sensor tips. Full-length KCTD1 protein was titrated up to 20 µM to assess binding. (**C**) KCTD1 a.a 28-257 shows no binding. (**D**) Full-length KCTD1 shows dose depending binding to TFAP2A with *K*_D_ = 380 nM by BLI. Protein was buffered in 25 mM HEPES pH 7.5, 150 mM NaCl, 0.05% Tween 20, 0.5 mM TCEP.

### Iodide binding to KCTD1 CTD affects protein stability and TFAP2 binding

Electron density maps indicated the presence of sodium and iodide ions in the central channel formed by the five CTD βD strands (Figures 3A and S2). By optimizing the crystallization conditions, we were able to increase the occupancy of two iodides to unity. Their identity was further confirmed by a diffraction dataset collected at 7000 eV that revealed a peak anomalous signal of 15σ. Interestingly, the sodium ions were coordinated by the backbone carbonyls and side chain hydroxyls of a βD Ser-X-Gly-X-Gly motif that was reminiscent of the same groups in the Thr-X-Gly-X-Gly selectivity sequence of the pore loop domain in the Kv channel family (Figure S2).

**Figure 3.**
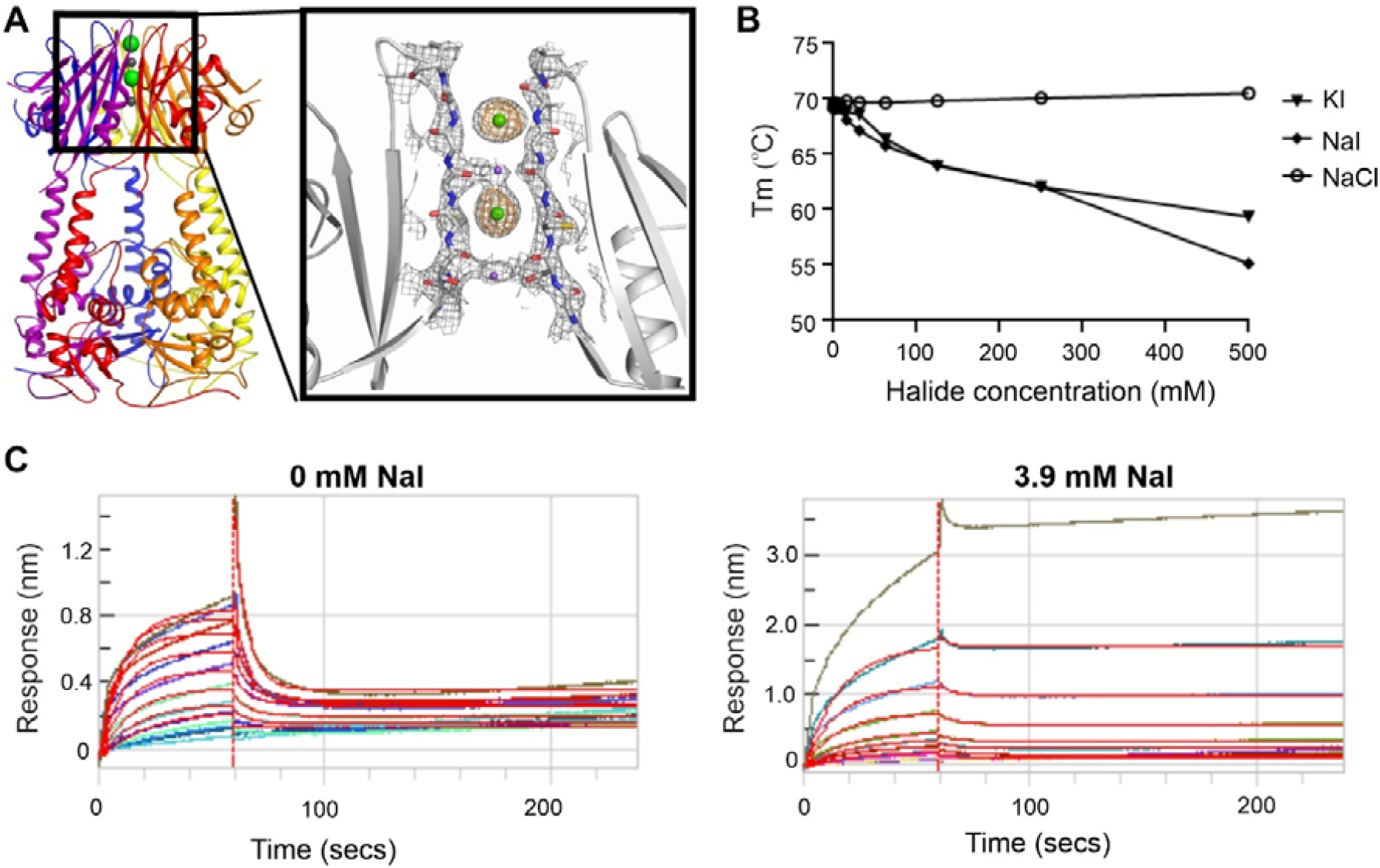
Iodide binding to KCTD1 CTD affects protein stability and TFAP2 binding. (**A**) KCTD1 pentamer forms an ‘ion pore’ in the CTD occupied by two sodium and two iodide ions (purple and green spheres, respectively). Inset slice shows 2Fo-Fc electron density map (gray) contoured at 1.0 σ overlaid with iodide-specific anomalous map (brown) contoured at 5.0 σ. (**B**) Apparent melting temperature of full-length KCTD1 at different halide ion salt concentrations. Protein was buffered in 25 mM HEPES pH 7.5, 0.05% Tween-20 plus indicated halide salts and SYPRO orange dye. Dilutions from 500 mM salt were mixed with NaH_2_PO_4_ pH 7.5 to maintain constant ionic strength. (**C**) BLI measurements of KCTD1 binding to immobilised TFAP2A peptide. Full-length KCTD1 protein was titrated up to 20 µM to assess binding buffered in 25 mM HEPES pH 7.5, 150 mM NaCl, 440 mM NaH2PO4 pH 7.5, 0.05% Tween 20 and indicated NaI concentrations. See also Figures S2-S3.

We investigated whether the bound ions were relevant to the structure and function of KCTD1. Sodium and potassium iodide salts were found to be destabilising as shown by a reduction in the apparent melting temperature of the full-length protein (Figures 3B and S3A). This effect was specific to iodide as the melting temperature was unaffected by sodium chloride concentration irrespective of conditions of constant (Figure 3B) or variable ionic strength (Figure S3A). We extended these experiments to the isolated BTB and CTD domains and observed that the destabilisation induced by iodide occurred only with the CTD consistent with the ion binding site, whereas the BTB domain was stabilised (Figures S3B-C). Finally, we found that even low millimolar concentrations of sodium iodide were sufficient to alter the binding of KCTD1 to TFAP2A and to limit the dissociation of the peptide from the bound complex (Figure 3C). These results show that sodium iodide binding to the CTD affects both the stability and ligand interactions of KCTD1.

### KCTD1 CTD shares structural homology with GFRP as well as KCTD5 and KCTD12

The CTD structure comprises a five-stranded antiparallel β-sheet (βA-βE) interrupted by two α-helices (αA, αC), and a 3_10_-helix (αB)(Figures 1C-D and 4A). Prior to structure determination, the KCTD1 CTD showed 80% sequence identity to its paralogue KCTD15, but had no other obvious matches by sequence homology. The structure helps to define a 40-residue insertion (βB-αA-βC-αB) relative to KCTD5 that enables its structural conservation to be recognized (Figures 1C-D). In the pentameric assembly, these structural elements lie on the outermost face of the β-propeller-like fold, where they potentially form a novel interface for binding partners, as observed for the prototypical KCTD5^6,7^.

A search for structural homologues of the KCTD1 CTD using the DALI server^23^ identified GTP Cyclohydrolase I Feedback Regulatory Protein (GFRP) as an unexpected match outside of the KCTD family (RMSD = 2.3 Å over 83 Cα atoms, Figures 4B-C), a protein for which there is virtually no sequence similarity (Figure 4D). Moreover, the GFRP protein assembles into a closed pentameric complex nearly identical to that of the CTD of KCTD1 (Figures 4A-B). To our surprise, the KCTD12 CTD domain was another close match (RMSD = 2.2 Å over 83 Cα atoms) revealing far wider conservation within the KCTD family than previously thought (Figure S4). It should be noted, however, that the GFRP, KCTD5 and KCTD12 structures all show ‘open’ solvent-filled central channels in contrast to the tightly restricted ion-bound channel of KCTD1. The clade F proteins KCTD8/12/16 are also distinguished from other KCTD proteins by having large flexible linkers between their BTB and CTD domains (Figure S4).

**Figure 4.**
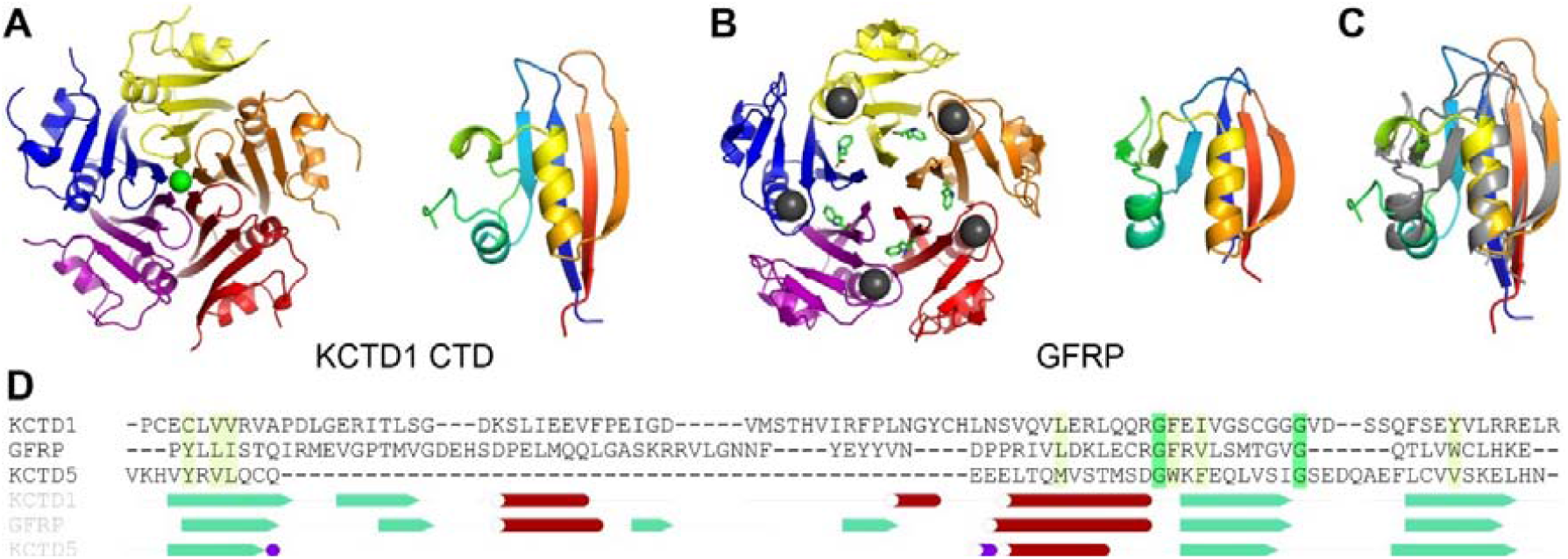
KCTD1 CTD shares structural homology with GFRP. (**A**) KCTD1 CTD pentamer (left) and subunit topology by rainbow colours (right). (**B**) Human GFRP pentamer (PDB 7AL9, left) and chain topology (right). Bound phenylalanine and potassium ions are shown as green sticks and gray spheres, respectively. (**C**) Superposition of KCTD1 CTD (rainbow) and GFRP (gray) chains. (**D**) Structure-based sequence alignment. See also Figure S4.

Overall, the elucidation of the full-length KCTD1 structure uncovers novel and important features of the BTB and CTD domains that contribute to its oligomeric assembly and protein-ligand interactions.

## DISCUSSION

This work unveils novel features of the KCTD1 pentamer, as well as unanticipated general lessons for the KCTD family. The BTB domain is the most characterized region of the KCTDs, but is habitually studied using truncated expression constructs. The identified N-terminal extension of the KCTD1 BTB domain presents a strikingly parallel to the β1-strand of the dimeric BTB protein class. In dimeric BTBs, the N-terminal β1-strand establishes domain-swap interactions that stabilize the dimer interface^24^, as well as a binding surface for transcriptional repressors^25^. In an analogous arrangement, the N-terminal region of KCTD1 forms cyclic domain-swap interactions to stabilize the BTB pentamer, as well as a critical region for the binding of TFAP2A. This explains how the N-terminal P20S mutation identified in scalp-ear-nipple syndrome disrupts both the thermostability of KCTD1 and its TFAP2A interaction^19^. We anticipate that this functionally important BTB extension will be conserved in other KCTD family members and advise that future studies incorporate it at construct design.

The BTB and CTD domains are tethered by a helix that may allow functional interplay between the two pentameric rings, as evidenced by E3 ligases such as KCTD5^6,7^. We note that the CTD of KCTD5 displays a relatively ‘open’ solvent-filled central channel, whereas the channel of KCTD1 is restricted to bind to alternating sodium and iodide ions, the most unusual feature of the structure. Similar to the pre-BTB region, ion concentration was found to modulate the TFAP2A interaction suggestive of potential BTB-CTD cross-talk. Despite co-crystallization, we were unable to identify bound TFAP2A peptide in our structure to shed further insight into this relationship. This interaction may be defined in future by cryo-EM studies using the full-length TFAP2A protein. The ion-binding of KCTD1 is noteworthy as the KCTD1/TFAP2A transcriptional axis regulates the reabsorption of salts in the kidney^22^.

The CTD structure also exposes fold similarities between disparate clades of the KCTD family that are otherwise difficult to discern from their sequences alone. Particularly striking is the structural conservation of KCTD1 (clade A) and KCTD12 (clade F), and their similar insertion relative to the CTD of the prototypical KCTD5 (clade E). These observations strengthen the phylogenetic linkage between the prototypical KCTD family members acting as E3 ligase adaptors and those with alternative functions^1^. The surprising similarity of the CTDs to GFRP also raises the possibility that these proteins may have arisen from a common ancestor. GFRP acts as a metabolic sensor that also depends on its pentameric structure to bind to GTP cyclohydrolase I (GCH1) as an allosteric inhibitor or activator dependent on the availability of cofactors tetrahydrobiopterin or phenylalanine, respectively^26-28^.

A common thread throughout this work is the importance of oligomerization for KCTD1 function, as evidenced by the contributions of the pre-BTB extension, ion-binding CTD and dominant-negative mutations in scalp-ear-nipple syndrome. Our work sheds further light on the structure-function relationship of KCTD proteins, and lays a structural foundation for understanding the non-E3 KCTDs.

## METHODS

### Cloning

Constructs were prepared using ligation-independent cloning. Full-length human KCTD1 (Uniprot Q719H9, residues 1-257) was inserted into the pNIC-NHStIIT vector which provides N-terminal hexahistidine and tandem Streptag II tags followed by a tobacco etch virus (TEV) cleavage site. The didomain construct of KCTD1 containing residues 28-257 was cloned into the pCDF-LIC vector which provides an N-terminal hexahistidine tag and TEV cleavage site. The KCTD1 BTB domain (residues 26-132) was similarly cloned into the pCDF-LIC vector, while the KCTD1 CTD domain (residues 132-257) was cloned into the pNIC28-Bsa4 vector, which also provides an N-terminal hexahistidine tag and TEV cleavage site

### Expression

Constructs were transformed into BL21(DE3)-R3-pRARE2 cells. Expression was performed in Luria-Bertani (LB) broth supplemented with appropriate antibiotics (kanamycin and chloramphenicol for the pNIC28-Bsa4 and pNIC-NHStIIT vector constructs, and spectinomycin and chloramphenicol for the pCDF-LIC vector constructs). Cultures were grown at 37°C with shaking at 180 RPM to an OD_600_ value of 0.6-0.8 and then cooled to 18°C before induction overnight by addition of 0.3 mM isopropyl β-D-1-thiogalactopyranoside (IPTG). Cells were harvested by centifugation at 5000 g for 10 mins.

### Purification

Cell pellets were resuspended in binding buffer (50 mM HEPES pH 7.5, 500 mM NaCl, 5 mM Imidazole, 5 % glycerol) supplemented with Protease Inhibitor Cocktail Set III, EDTA-Free and 0.5 mM tris(2-carboxyethyl)phosphine (TCEP). Cells were then lysed by sonication performed on ice. Lysates were cleared by centrifugation at 4°C in a JA25.5 rotor for 45 minutes at 36,200 g. Hexahistidine-tagged KCTD1 proteins were purified from the supernatant using nickel-sepharose resin. Captured proteins were washed with binding buffer supplemented with 30 mM imidazole and then eluted stepwise with binding buffer supplemented with 100, 150, 250 or 300 mM imidazole. The flow through, wash and elution fractions were analysed by SDS PAGE to identify KCTD1-containing fractions, which were then pooled and cleaved overnight with TEV protease. A final clean up step was performed using size exclusion chromatography using a S200 superdex column buffered in 50 mM HEPES pH 7.5, 300 mM NaCl, 5% glycerol, 1 mM TCEP.

### Crystallisation

Proteins were concentrated to 10 mg/mL using vivaspin 10 MWCO concentrators (GE healthcare). Crystallisation screens were performed by sitting drop vapour diffusion using 3 Lens Crystallisation Plates (SWISSCI, UK) at 4°C and 20°C and explored six different sparse matrix precipitant screens (LFS, JCSG, HCS, HIN, BCS and SaltRx). Three 150 nL sitting droplets per well were prepared by mixing protein and precipitant in ratios of 2:1, 1:1 and 1:2. Didomain KCTD1_Δ__N27_ containing residues 28-257 was buffered in 50 mM HEPES pH 7.5, 300 mM NaCl, 5 % glycerol, 1 mM TCEP. Diffracting crystals grew at 20°C in drops mixing 50 nL protein with 100 nL of a mother liquor containing 12% PEG3350, 0.1 M bis-tris-propane pH 7.8, 0.2 M sodium iodide, 10% ethylene glycol. Mounted crystals were cryoprotected in acetonitrile and vitrified in liquid nitrogen. Full-length KCTD1 was buffered in 50 mM HEPES pH 7.5, 300 mM NaCl, 200mM NaI, 0.5 mM TCEP and mixed 1:1 with TFAP2A (AP-2α) peptide (NADFQPPYFPPPYQ, transactivation domain residues 50–63, Uniprot P05549-1,). Diffracting crystals grew at 4°C in drops mixing 100 nL protein with 50 nL of a mother liquor containing 0.1 M SPG pH 6.0 (mix ratio 2:7:7 succinate/sodium dihydrogen phosphate/glycine), 60.0% MPD (2-methyl-2,4-pentanediol). Mounted crystals were cryoprotected in mother liquor plus 25% ethylene glycol and vitrified in liquid nitrogen.

### Diffraction data collection and structure refinement

Diffraction data were collected at 100 K on Diamond Light Source beamline I04 (KCTD1 didomain) or I24 (KCTD1 full length). Data were indexed and integrated using XDS and scaled using AIMLESS. The structures were solved by molecular replacement in PHASER.

The pentameric KCTD1 BTB domain structure PDB 5BXB was used as an initial search model for KCTD1_Δ__N27_ and this structure was subsequently used as a search model for the full-length KCTD1 structure. Initial electron density maps for KCTD1_Δ__N27_ indicated the presence of ions in the central tunnel of the CTD1, consistent with partial occupancy of sodium and iodide sequestered from the crystallization solution. By optimizing the crystallization conditions, we were able to increase the occupancy of two Iodides in the central tunnel to unity. To further confirm the identity of the ions, a dataset was collected at 7000 eV. The Iodides exhibited a peak anomalous signal of 15σ (Figure 3A). Manual model building was performed in COOT and refinement completed using PHENIX.REFINE. The refined structures were validated with MolProbity and the atomic coordinate files deposited in the Protein Data Bank. Structure figures were generated using PyMOL

### Biolayer interferometry

Biolayer interferometry (BLI) experiments were performed on an Octet RED384 instrument (FortéBio). Experiments were performed in a buffer containing 25 mM HEPES pH 7.5, 150 mM NaCl, 0.05% (v/v) Tween 20, 0.5 mM TCEP unless stated otherwise. Biotinylated TFAP2A (AP-2α) peptide (sequence: Biotin-NADFQPPYFPPPYQ, Uniprot P05549-1 residues 50–63) was immobilised onto streptavidin-coated fibre optic sensor tips, while ‘free’ reference sensors were used without immobilised peptide. Sensor tips were sequentially dipped into their respective protein samples from low to high concentrations for association phases and then into buffer for dissociation phases. Data were analysed using the FortéBio Data Analysis software.

### Differential scanning fluorimetry

Purified proteins in the indicated buffers were mixed with SYPRO Orange dye (1:1000 dilution). Samples were heated from 25°C to 95°C in a Mx3005P real-time PCR instrument (Stratagene). Fluorescence was monitored with excitation and emission filters set to 465 and 590 nm, respectively. Data were exported to GraphPad Prism and fitted to the Boltzmann equation to calculate apparent *T*_m_ values as described previously. Full-length and BTB domain constructs were assayed at 2 µM protein concentration, whereas the KCTD1 CTD construct was assayed at 100 µM protein concentration due to low signal.

## ACKNOWLEDGEMENTS

The authors would like to thank Diamond Light Source for beamtime (proposals mx15433 and mx19301), as well as the staff of beamlines I04 and I24 for assistance with crystal testing and data collection. ANB acknowledges funding from the Innovative Medicines Initiative 2 Joint Undertaking (JU) under grant agreement number 875510 (EUbOPEN). JU receives support from the European Union’s Horizon 2020 research and innovation programme and EFPIA and Ontario Institute for Cancer Research, Royal Institution for the Advancement of Learning McGill University, Kungliga Tekniska Hoegskolan, Diamond Light Source. This research was also funded in part by a Wellcome strategic award (106169/ZZ14/Z).

## AUTHOR CONTRIBUTIONS

Conceptualisation DMP, ANB; investigation DMP, JCB, AEH, CEM, WR; supervision DMP, ANB; manuscript preparation DMP, ANB; funding acquisition ANB. All authors reviewed the submitted manuscript.

## Data Availability

The coordinates and structure factors for the crystal structures reported in this article have been deposited in the PDB with accession codes 9FOI (full length KCTD1) and 6S4L (KCTD1_Δ__N27_).

## SUPPLEMENTAL INFORMATION

Document S1. Table S1 and Figures S1–S4

## References

1. Skoblov, M., Marakhonov, A., Marakasova, E., Guskova, A., Chandhoke, V., Birerdinc, A., and Baranova, A. (2013). Protein partners of KCTD proteins provide insights about their functional roles in cell differentiation and vertebrate development. Bioessays 35, 586–596. 10.1002/bies.201300002.

2. Balasco, N., Pirone, L., Smaldone, G., Di Gaetano, S., Esposito, L., Pedone, E.M., and Vitagliano, L. (2014). Molecular recognition of Cullin3 by KCTDs: insights from experimental and computational investigations. Biochim Biophys Acta 1844, 1289–1298. 10.1016/j.bbapap.2014.04.006.

3. Ji, A.X., Chu, A., Nielsen, T.K., Benlekbir, S., Rubinstein, J.L., and Prive, G.G. (2016). Structural Insights into KCTD Protein Assembly and Cullin3 Recognition. J Mol Biol 428, 92–107. 10.1016/j.jmb.2015.08.019.

4. Pinkas, D.M., Sanvitale, C.E., Bufton, J.C., Sorrell, F.J., Solcan, N., Chalk, R., Doutch, J., and Bullock, A.N. (2017). Structural complexity in the KCTD family of Cullin3-dependent E3 ubiquitin ligases. Biochem J 474, 3747–3761. 10.1042/BCJ20170527.

5. Dementieva, I.S., Tereshko, V., McCrossan, Z.A., Solomaha, E., Araki, D., Xu, C., Grigorieff, N., and Goldstein, S.A. (2009). Pentameric assembly of potassium channel tetramerization domain-containing protein 5. J Mol Biol 387, 175–191. 10.1016/j.jmb.2009.01.030.

6. Jiang, W., Wang, W., Kong, Y., and Zheng, S. (2023). Structural basis for the ubiquitination of G protein betagamma subunits by KCTD5/Cullin3 E3 ligase. Sci Adv 9, eadg8369. 10.1126/sciadv.adg8369.

7. Nguyen, D.M., Rath, D.H., Devost, D., Petrin, D., Rizk, R., Ji, A.X., Narayanan, N., Yong, D., Zhai, A., Kuntz, D.A., et al. (2024). Structure and dynamics of a pentameric KCTD5/CUL3/Gbetagamma E3 ubiquitin ligase complex. Proc Natl Acad Sci U S A 121, e2315018121. 10.1073/pnas.2315018121.

8. Brockmann, M., Blomen, V.A., Nieuwenhuis, J., Stickel, E., Raaben, M., Bleijerveld, O.B., Altelaar, A.F.M., Jae, L.T., and Brummelkamp, T.R. (2017). Genetic wiring maps of single-cell protein states reveal an off-switch for GPCR signalling. Nature 546, 307–311. 10.1038/nature22376.

9. Sloan, D.C., Cryan, C.E., and Muntean, B.S. (2023). Multiple potassium channel tetramerization domain (KCTD) family members interact with Gbetagamma, with effects on cAMP signaling. J Biol Chem 299, 102924. 10.1016/j.jbc.2023.102924.

10. Smaldone, G., Pirone, L., Balasco, N., Di Gaetano, S., Pedone, E.M., and Vitagliano, L. (2015). Cullin 3 Recognition Is Not a Universal Property among KCTD Proteins. PLoS One 10, e0126808. 10.1371/journal.pone.0126808.

11. Zheng, S., Abreu, N., Levitz, J., and Kruse, A.C. (2019). Structural basis for KCTD-mediated rapid desensitization of GABAB signalling. Nature 567, 127–131. 10.1038/s41586-019-0990-0.

12. Zuo, H., Glaaser, I., Zhao, Y., Kurinov, I., Mosyak, L., Wang, H., Liu, J., Park, J., Frangaj, A., Sturchler, E., et al. (2019). Structural basis for auxiliary subunit KCTD16 regulation of the GABA(B) receptor. Proc Natl Acad Sci U S A 116, 8370–8379. 10.1073/pnas.1903024116.

13. Turecek, R., Schwenk, J., Fritzius, T., Ivankova, K., Zolles, G., Adelfinger, L., Jacquier, V., Besseyrias, V., Gassmann, M., Schulte, U., et al. (2014). Auxiliary GABAB receptor subunits uncouple G protein betagamma subunits from effector channels to induce desensitization. Neuron 82, 1032–1044. 10.1016/j.neuron.2014.04.015.

14. Correale, S., Esposito, C., Pirone, L., Vitagliano, L., Di Gaetano, S., and Pedone, E. (2013). A biophysical characterization of the folded domains of KCTD12: insights into interaction with the GABAB2 receptor. J Mol Recognit 26, 488–495. 10.1002/jmr.2291.

15. Hu, L., Chen, L., Yang, L., Ye, Z., Huang, W., Li, X., Liu, Q., Qiu, J., and Ding, X. (2020). KCTD1 mutants in scalp⍰ear⍰nipple syndrome and AP⍰2alpha P59A in Char syndrome reciprocally abrogate their interactions, but can regulate Wnt/beta⍰catenin signaling. Mol Med Rep 22, 3895–3903. 10.3892/mmr.2020.11457.

16. Zarelli, V.E., and Dawid, I.B. (2013). Inhibition of neural crest formation by Kctd15 involves regulation of transcription factor AP-2. Proc Natl Acad Sci U S A 110, 2870–2875. 10.1073/pnas.1300203110.

17. Miller, K.A., Cruz Walma, D.A., Pinkas, D.M., Tooze, R.S., Bufton, J.C., Richardson, W., Manning, C.E., Hunt, A.E., Cros, J., Hartill, V., et al. (2024). BTB domain mutations perturbing KCTD15 oligomerisation cause a distinctive frontonasal dysplasia syndrome. J Med Genet 61, 490–501. 10.1136/jmg-2023-109531.

18. Raymundo, J.R., Zhang, H., Smaldone, G., Zhu, W., Daly, K.E., Glennon, B.J., Pecoraro, G., Salvatore, M., Devine, W.A., Lo, C.W., et al. (2023). KCTD1/KCTD15 complexes control ectodermal and neural crest cell functions, and their impairment causes aplasia cutis. J Clin Invest 134. 10.1172/JCI174138.

19. Smaldone, G., Balasco, N., Pirone, L., Caruso, D., Di Gaetano, S., Pedone, E.M., and Vitagliano, L. (2019). Molecular basis of the scalp-ear-nipple syndrome unraveled by the characterization of disease-causing KCTD1 mutants. Sci Rep 9, 10519. 10.1038/s41598-019-46911-4.

20. Milunsky, J.M., Maher, T.A., Zhao, G., Roberts, A.E., Stalker, H.J., Zori, R.T., Burch, M.N., Clemens, M., Mulliken, J.B., Smith, R., and Lin, A.E. (2008). TFAP2A mutations result in branchio-oculo-facial syndrome. Am J Hum Genet 82, 1171–1177. 10.1016/j.ajhg.2008.03.005.

21. Marneros, A.G., Beck, A.E., Turner, E.H., McMillin, M.J., Edwards, M.J., Field, M., de Macena Sobreira, N.L., Perez, A.B., Fortes, J.A., Lampe, A.K., et al. (2013). Mutations in KCTD1 cause scalp-ear-nipple syndrome. Am J Hum Genet 92, 621–626. 10.1016/j.ajhg.2013.03.002.

22. Marneros, A.G. (2020). AP-2beta/KCTD1 Control Distal Nephron Differentiation and Protect against Renal Fibrosis. Dev Cell 54, 348–366 e345. 10.1016/j.devcel.2020.05.026.

23. Holm, L., and Rosenstrom, P. (2010). Dali server: conservation mapping in 3D. Nucleic Acids Res 38, W545–549. 10.1093/nar/gkq366.

24. Ahmad, K.F., Engel, C.K., and Prive, G.G. (1998). Crystal structure of the BTB domain from PLZF. Proc Natl Acad Sci U S A 95, 12123–12128. 10.1073/pnas.95.21.12123.

25. Ahmad, K.F., Melnick, A., Lax, S., Bouchard, D., Liu, J., Kiang, C.L., Mayer, S., Takahashi, S., Licht, J.D., and Prive, G.G. (2003). Mechanism of SMRT corepressor recruitment by the BCL6 BTB domain. Mol Cell 12, 1551–1564. 10.1016/s1097-2765(03)00454-4.

26. Bader, G., Schiffmann, S., Herrmann, A., Fischer, M., Gutlich, M., Auerbach, G., Ploom, T., Bacher, A., Huber, R., and Lemm, T. (2001). Crystal structure of rat GTP cyclohydrolase I feedback regulatory protein, GFRP. J Mol Biol 312, 1051–1057. 10.1006/jmbi.2001.5011.

27. Maita, N., Okada, K., Hatakeyama, K., and Hakoshima, T. (2002). Crystal structure of the stimulatory complex of GTP cyclohydrolase I and its feedback regulatory protein GFRP. Proc Natl Acad Sci U S A 99, 1212–1217. 10.1073/pnas.022646999.

28. Ebenhoch, R., Prinz, S., Kaltwasser, S., Mills, D.J., Meinecke, R., Rubbelke, M., Reinert, D., Bauer, M., Weixler, L., Zeeb, M., et al. (2020). A hybrid approach reveals the allosteric regulation of GTP cyclohydrolase I. Proc Natl Acad Sci U S A 117, 31838–31849. 10.1073/pnas.2013473117.

