## Supplemental Table S1 and Figures S1-4 for "A BTB extension and ion-binding domain contribute to the pentameric structure and TFAP2A binding of KCTD1"

**Table S1. Diffraction data collection and structure refinement statistics**

| | KCTD1 full length<br>PDB ID 9FOI | $KCTD1_{\Delta N27}$<br>PDB ID 6S4L |
|---|---|---|
| Wavelength (Å) | 0.9686 | 1.771 |
| Resolution range (Å) | 114.84 - 2.71 (2.85 - 2.71) | 67.41 - 2.8 (2.9 - 2.8) |
| Space group | P 1 21 1 | P 1 21 1 |
| Unit cell a, b, c (Å)<br>α, β, γ (°) | 67.558 96.634 115.995 90<br>98.088 90 | 67.454 95.87 104.343 90<br>92.158 90 |
| Total reflections | 277262 (27857) | 234273 (24285) |
| Unique reflections | 40265 (3965) | 31509 (3082) |
| Multiplicity | 6.9 (6.9) | 7.4 (7.9) |
| Completeness (%) | 99.79 (98.85) | 95.88 (94.98) |
| Mean I/sigma(I) | 7.01 (1.1) | 7.56 (1.16) |
| Wilson B-factor | 57.05 | 53.69 |
| R-merge | 0.213 (1.910) | 0.2596 (2.066) |
| R-meas | 0.251 (2.253) | 0.2796 (2.213) |
| R-pim | 0.132 (1.182) | 0.1023 (0.7886) |
| CC1/2 | 0.993 (0.428) | 0.99 (0.529) |
| CC* | 0.998 (0.755) | 0.998 (0.832) |
| Reflections used in refinement | 40308 (3970) | 31497 (3082) |
| Reflections used for R-free | 1954 (186) | 1481 (159) |
| R-work | 0.206 (0.3104) | 0.2466 (0.3580) |
| R-free | 0.245 (0.3708) | 0.2759 (0.3883) |
| Number of non-hydrogen atoms | 8500 | 8534 |
| macromolecules | 8352 | 8468 |
| ligands | 85 | 4 |
| solvent | 63 | 62 |
| Protein residues | 1048 | 1060 |
| RMSD (bonds, Å) | 0.0046 | 0.003 |
| RMSD (angles, °) | 0.70 | 0.90 |
| Ramachandran favored (%) | 94.26 | 92.60 |
| Ramachandran allowed (%) | 5.74 | 7.40 |
| Ramachandran outliers (%) | 0.00 | 0.00 |
| Rotamer outliers (%) | 0.44 | 2.39 |
| Clashscore | 2.13 | 6.69 |
| Average B-factor ($Å^2$) | 64.8 | 53.90 |
| macromolecules | 64.9 | 54.08 |
| ligands | 62.3 | 53.59 |
| solvent | 51.0 | 30.15 |

Statistics for the highest-resolution shell are shown in parentheses.

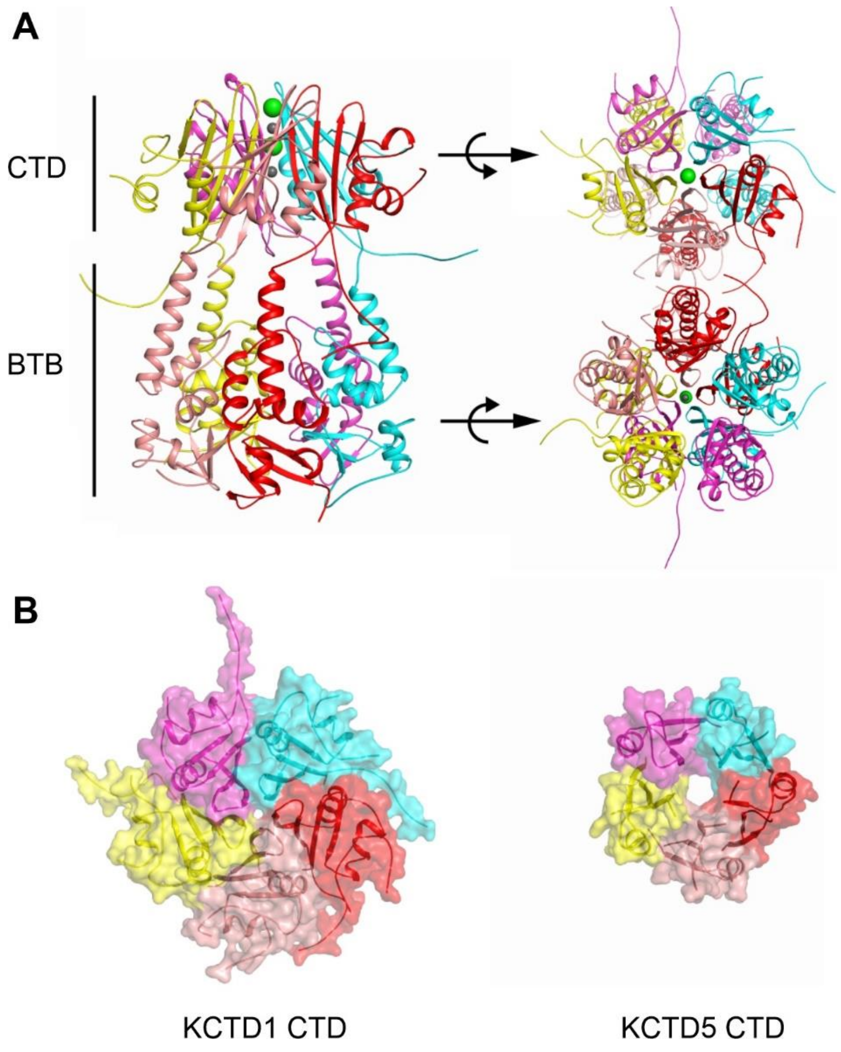


**Figure S1. Structure of KCTD1$_{\Delta N27}$**

(**A**) Ribbon diagram of the pentameric structure of KCTD1$_{\Delta N27}$. Regions spanning the BTB and C-terminal domains (CTD) are indicated. Different protomers within the pentamer are distinguished by different colours. Sodium and iodide ions located in the CTD are shown as gray and green spheres, respectively. Right panel shows 90° rotations to show views from the top and bottom perspective of the image, respectively. (**B**) Transparent surface representations of the KCTD1 and KCTD5 CTD pentamers. The central channel in KCTD5 (right) is more open than the ion-bound channel of KCTD1 (left).

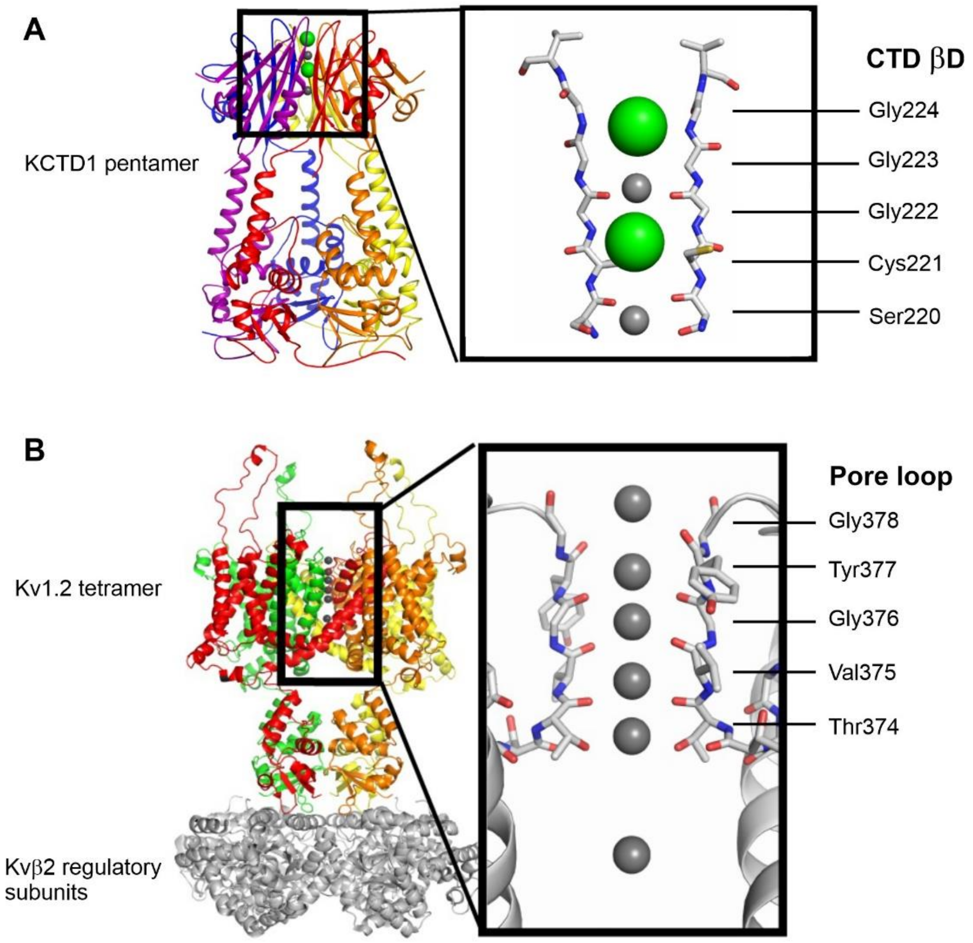


**Figure S2. Backbone carbonyls form the ion pore in KCTD1**

(**A**) Ribbon diagram of the pentameric KCTD1 structure. Inset shows a slice through the 'ion pore' (not membrane spanning) in the CTD occupied by two sodium and two iodide ions (gray and green spheres, respectively). The CTD pentamer presents five βD strands that line the pore with their backbone carbonyls. Each strand has a Ser-X-Gly-X-Gly motif that is reminiscent of the Thr-X-Gly-X-Gly motif of pore loop domain (P-domain) selectivity sequence in the Kv channel family. The backbone carbonyls and side chain hydroxyl from the Ser-X-Gly are observed to coordinate the sodium ions. (**B**) Ribbon diagram of the rat Kv1.2 shaker potassium channel (PDB 3LUT). The four subunits of the tetramer are indicated by different colours. The associated Kvβ2 regulatory subunits are coloured gray. Inset shows a slice through the ion pore with potassium ions shown as gray spheres. Residues in the Thr-X-Gly-X-Gly motif of the pore loop domain (P-domain) selectivity sequence are labelled.

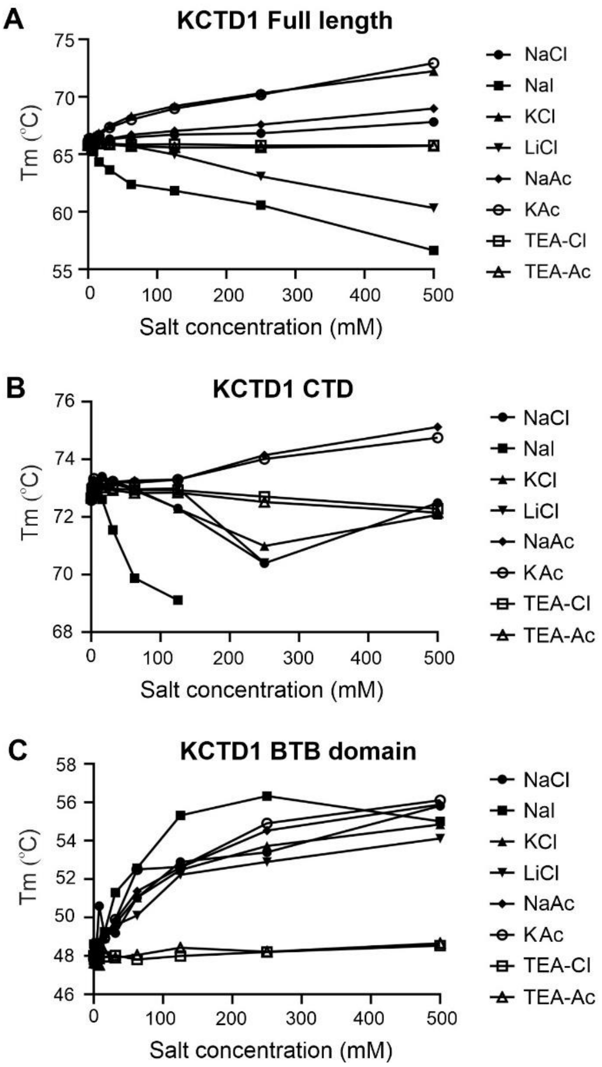


**Figure S3. Iodide destabilizes KCTD1 CTD folding**

DSF experiments showing apparent melting temperature of (**A**) full-length KCTD1, as well as the isolated (**B**) CTD or (**C**) BTB domains under different salt conditions and concentrations without adjustments for ionic strength. Sodium iodide destabilises the CTD as well as the full-length protein, whereas the BTB domain is stabilized. Protein was buffered in 25 mM HEPES pH 7.5, 0.05% Tween-20 plus indicated salts and SYPRO orange dye. Full length and BTB domain constructs were assayed at 2 µM protein concentration, whereas the KCTD1 CTD construct was assayed at 100 µM protein concentration due to low fluorescent signal. Signal changes for the KCTD1 CTD were also not observable above a concentration of 125 mM NaI.

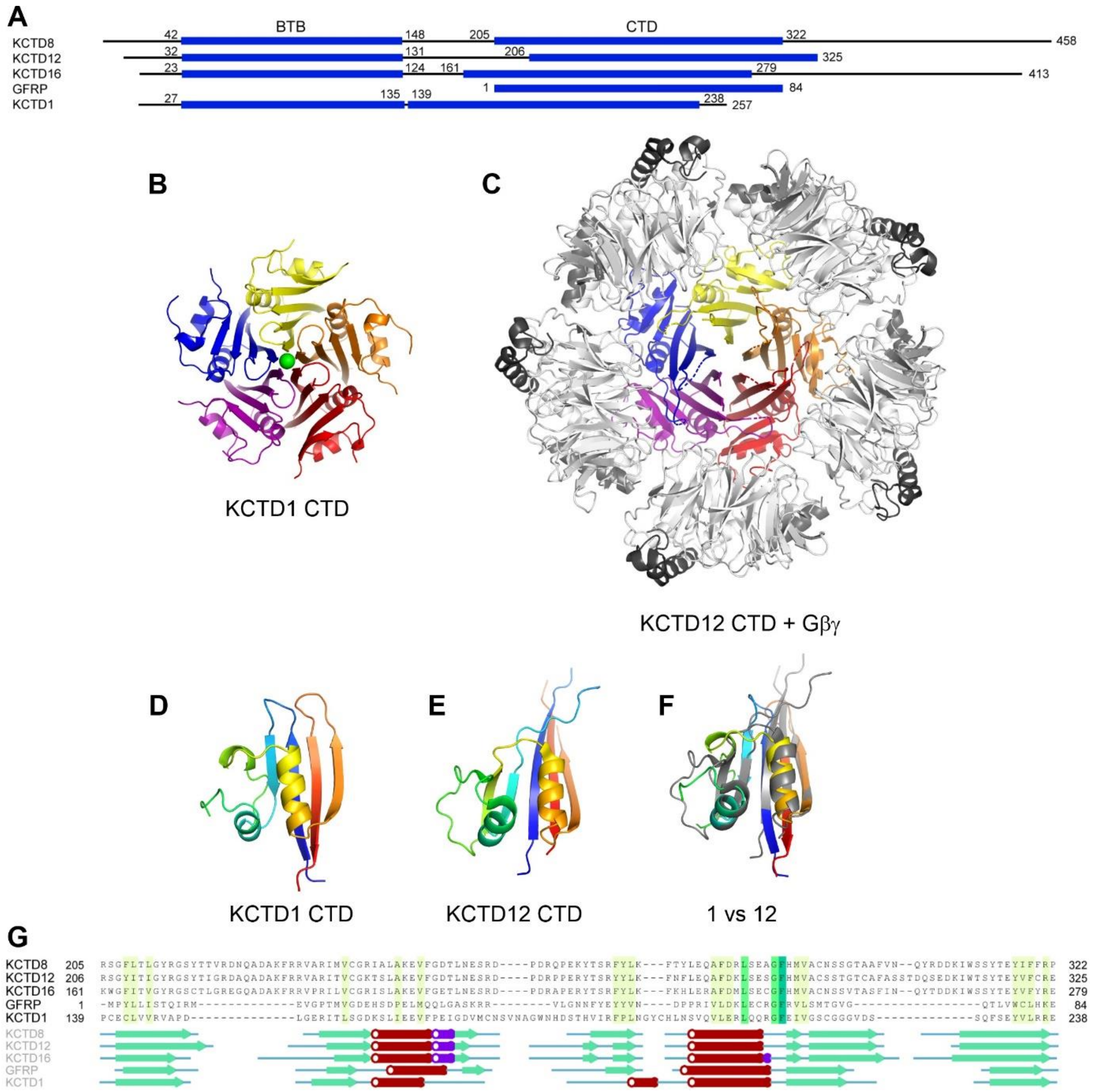


**Figure S4. KCTD1 CTD shows structural conservation with KCTD12**
(**A**) Domain organisation of KCTD1 and selected homologues. (**B**) KCTD1 CTD pentamer coloured by subunit. Bound iodide ion shown as green sphere. (**C**) KCTD12 CTD pentamer coloured by subunit bound to Gβ (light gray) and Gγ (dark gray) (PDB 6M8S). Subunit topology by rainbow colours for (**D**) KCTD1 CTD and (**E**) KCTD12 CTD. (**F**) Superposition of KCTD1 CTD (rainbow) and KCTD12 CTD (gray) chains. (G) Structure-based sequence alignment and secondary structure elements.
